# tacost: Testing and simulating the performance of acoustic tracking systems

**DOI:** 10.1101/2020.06.22.165308

**Authors:** Thejasvi Beleyur

## Abstract

`tacost` is a Python package to allow the testing of acoustic tracking systems. While many microphone array systems have been characterised analytically and experimentally – these are time-intensive methods. `tacost` provides a simulation based framework to rapidly assess the tracking behaviour of multiple array geometries, and the dissection of other relevant parameters. This paper explains briefly the design of the package and highlights two example use cases in which the tracking accuracy of different microphone geometries are characterised.

## Introduction

Acoustic tracking is a common method used to study vocalising animals such as birds, bats and cetaceans (Suzuki et al. 2017; Aubauer 1996; Møhl et al. 2000; Hügel et al. 2017; Holderied and Von Helversen 2003; Rhinehart et al., n.d.; Blumstein et al. 2011). Using acoustic tracking, biologists can detect the position of the animal and track it through space as it moves over time. The localisation accuracy of an acoustic tracking system depends on a variety of factors. There are *internal* factors such as microphone array geometry, signal processing routines, and the mathematical formulations used to localise sounds (time-of-arrival, time-of-arrival-difference, angle-of-arrival, power-steering). The *external* factors include aspects related to the actual signal itself, ie. signal-to-noise ratio, and spectro-temporal properties of the emitted sound (noise, linear/hyperbolic sweep) (Wahlberg 1999). While experiments and analytical modelling may be the definitive way to determine a tracking system’s end accuracy, simulations allow a quick and systematic method to estimate the source of tracking errors. `tacost` provides a flexible workflow to manipulate and study the effect of both internal and external factors. `tacost` generates audio files for source positions and array geometries specified by the user. This allows the user to analyse the efficacy of their tracking system’s baseline performance.

## Statement of need

Generating simulated audio for a set of source sounds, positions and a given array configuration is a relatively simple task. However, to this my knowledge, there are no publicly available, tested and documented packages for this task published to date. Codebases that are publicly available have the advantage of being used by a larger user-base and can thus benefit from bug discoveries much faster than in-house or individually written one-time use scripts. `tacost` provides a robust and well-documented software workflow (Taschuk and Wilson 2016) with user and developer friendly documentation hosted online. `tacost` contributes to the Python scientific ecosystem in the hope of promoting the growth of acoustics and bioacoustic research in open-source languages like Python. In particular, `tacost` will help researchers working in the field of acoustics and bio-acoustics (Framond-Bénard et al. 2020) plan and examine the behaviour of their acoustic tracking systems.

## Design

The design of `tacost` focusses on a reproducible and user-friendly method (Wilson et al. 2012) to generate WAV files that form the input for acoustic tracking softwares. Users may interact with `tacost` through custom-written Python scripts by calling it as a Python package with `import tacost` or in the ‘no-coding’ mode. The ‘no-coding’ mode is especially suitable for users unfamiliar with Python. The no-coding mode is based around a parameter file that is used to specify various parts of the WAV file to be created. Through the parameter file the user can specify the emitted sound, source positions, inter-sound-intervals, sampling rate and other relevant variables to customise the test scenario.

## Examples

The localisation accuracy of a microphone array may not be uniform over 3D space (Aubauer 1996; Wahlberg 1999). This accuracy is independent of the actual signal and recording conditions of the input data, but rather dependent on the array geometry and mathematical formulations used to record and calculate sound source position.

The accuracy of a few microphone array configurations has been characterised analytically (Aubauer 1996) and experimentally (Wahlberg 1999). While reflecting the system’s capabilities, analytical and experimental characterisations are often time-intensive. In contrast, simulation uncovers the intrinsic accuracy of an array relatively quickly through the use of audio files with simulated emission points spread across the recording volume of interest. `tacost` can be used to characterise the maximal localisation accuracy of an acoustic tracking system with novel array geometries and recording scenarios. In Example 1, I show how `tacost` can be used to verify known trends in localisation error with the tristar60, a commonly used array system. In Example 2, I show how `tacost` can be used to estimate the expected localisation error in a multi-microphone array with a novel and field-friendly geometry.

### 1. Localisation accuracy of the tristar60 system

The tristar60 array is a commonly used array geometry (Aubauer 1996; Holderied and Von Helversen 2003; Hügel et al. 2017; Lewanzik and Goerlitz 2018) with 4 microphones in a plane on an inverted T array. Three peripheral microphones are placed 120*◦* to each other at 60 cm distance from the central mic on the inverted T-array. A series of emission points spanning the upper right quadrant of the array were simulated. The emitted sound was set to a linear sweep. The output WAV files from `tacost` were run through the TOADSuite package (Goerlitz 2019; Stilz, n.d.; Hügel et al. 2017; Lewanzik and Goerlitz 2018), a software package that localises sounds using the time-of-arrival-differences across channels. Figure 1 shows the localisation accuracy map for the tristar60 microphone array. It can be seen that localisation error increases with increasing radial distance from the central microphone, and remains <7% of the radial distance.

**Figure 1:**
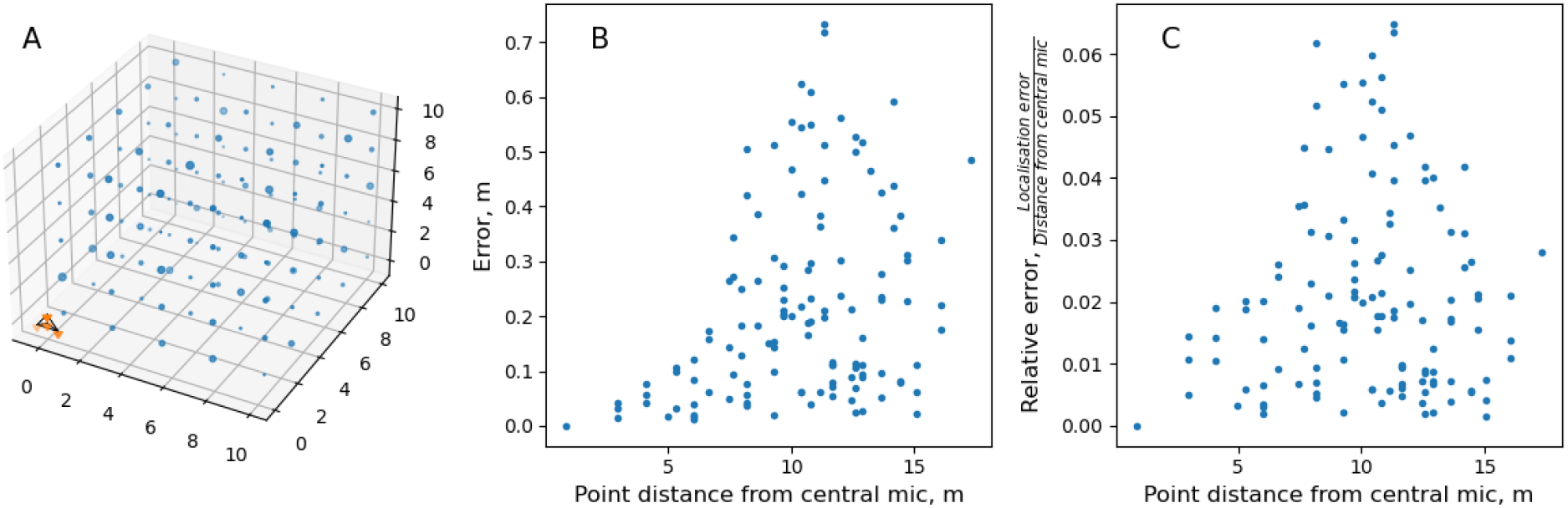
Accuracy of a sound source localisation based on time-of-arrival-differences with a tristar60 array. A) The tristar60 microphone array is placed at the origin of the coordinate system (bottom left, orange dots connected by black lines). The blue points are the simulated source positions which form a ‘calibration grid’. The size of each dot is proportional to the localisation error B) The localisation error increases with increasing radial distance of source from the central microphone.Each simulated point is shown as a dot, and the size of the dot is proportional to the tracking error. The error is the euclidean distance between the predicted and simulated source point.The errors range between 0-0.7m. C) The relative error of localisation. Even though absolute localisation error tends to increase with distance from array, all localisation happens with <7% relative error.

### 2. Localisation accuracy of a multi-microphone array in the field

While recording in the field, it may be difficult to use fixed arrays mounted on stands. Arrays on stands are difficult to carry and may also influence the behaviour of the animals being recorded. It is thus advantageous to use less obtrusive micorphone geometries, for instance by placing microphones are placed on pre-existing structures such as the walls of a cave or trees. These microphone geometries are field-friendly, but their localisation accuracy is hard to characterise analytically. `tacost` is an ideal tool to explore the tracking performance of such flexibly placed microphone arrays. Figure 2 shows the microphone array geometry and recording system described in (Batstone et al. 2019). In short, the array consisted of 11 microphones, 4 of them on a 120cm tristar, and the remaining 7 microphones attached to the walls of a cave. A series of sound emission points were created simulating points in the volume enclosed by the array. The points matched the volume echolocating bats flew within. The simulated sound was set to a linear sweep, which mimicked that of a bat call. The `tacost` output WAV files were analysed with the TOADSuite. The resulting accuracy map reveals that overall, the localisation error is between 7-30 centimetres for the given emission points. This corresponds to a maximum error of upto 30cm in tracking the position, and of upto 19% relative error. In contrast to the previous example highlighting the increase in tracking error with increasing source sound distance, these results show a somewhat different trend. The relative error is also much higher, and it may have to do with the positioning of the sound sources in the with reference to the array. The relative location of the sound source affects the tracking accuracy (Aubauer 1996).

**Figure 2:**
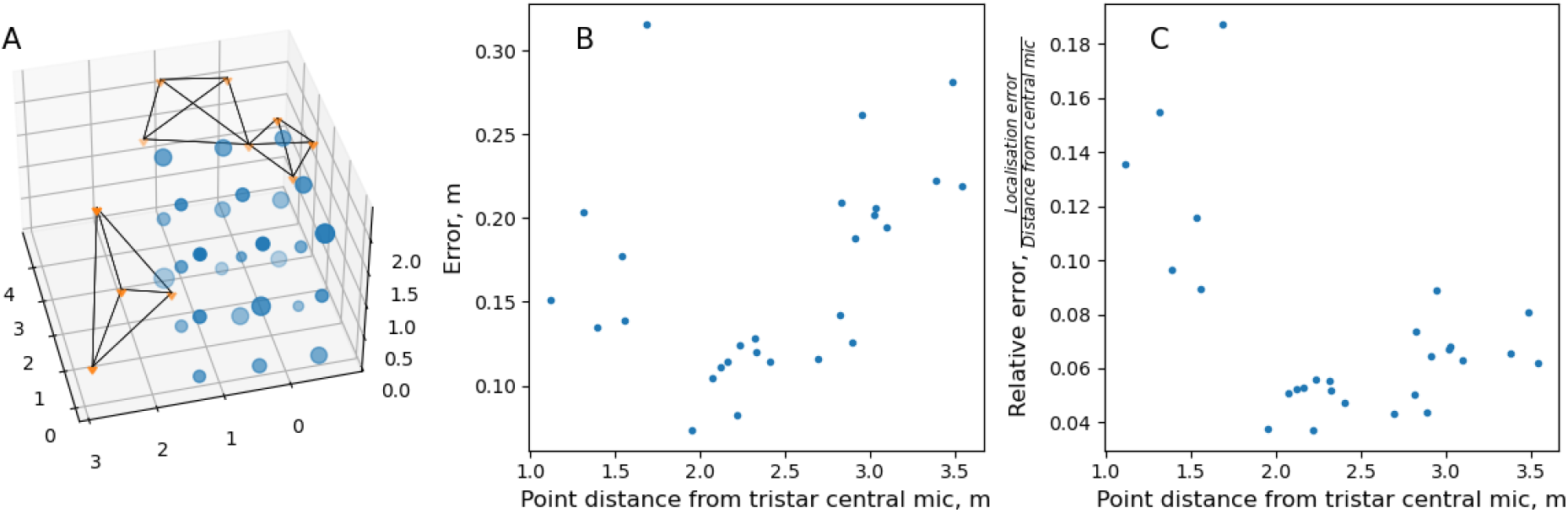
Localisation accuracy of a multi-microphone array in the field, localised with time-of-arrival-differences. A) The line-connected points (blue) represent the microphone array consisting of 11 microphones. Four microphones are in a tristar 120 array (left, tristar array with 120cm radial distance from central mic), and the remaining 7 mics are placed on the walls of the cave (top right, two quadirlateral outlines joined at a common vertex represent the 7 mics on the cave walls). The free-standing points (blue) are the simulated emission points which form a ‘calibration grid’ B) The distribution of localisation error. The error is the euclidean distance between the predicted and simulated point. Each simulated point is shown as a dot, and the size of the dot is proportional to the tracking error. The localisation error is between 0.07-0.32 m for the given points. C) The relative localisation error with reference to the central mic of the 120 cm tristar array microphone. The 95%ile bounds of tracking error lie between 3.7-16.6%, with a maximum of 18.8% error. Even points that are nearby seem to be localised with a higher relative error. This higher relative error may be the result of sound source position with reference to the microphone array.

## Future directions

`tacost` as it stands is currently written to implement a first-order assessment of a tracking system’s accuracy. The package has been primarily written keeping acoustic signals propagating through air where the velocity of sound is assumed to be constant. It may also be used to test tracking in radar or underwater sonar systems, contingent on how uniform the medium of wave propagation is over the distances being studied. As of version 0.1.0, straight line propagation of signals are simulated, without spherical spreading or atmospheric absorption implemented. Future releases may include such propagation losses. Another important aspect affecting all tracking systems is the directionality of the sensors (microphones) and emitted signals (animal vocalisations, calibration speakers). A common problem in acoustic tracking with bats and cetaceans is not being able to track animals because their echolocation calls can be very directional (Matsuta et al. 2013; Surlykke, Pedersen, and Jakobsen 2012; Koblitz et al. 2016). Implementing sensor and source sound directionality will help assessing how many microphones might be required to successfully track animals in their surroundings, and which array geometries are best able to do so.

## Acknowledgements

This work was supported by a doctoral fellowship from the German Academic Exchange Service (DAAD) and the International Max Planck Research School for Organismal Biology. I would like to thank Léna de Framond for generating the acoustic localisation output, Holger R Goerlitz for helpful comments on this manuscript and discussions on the topic of tracking, and the IT team atthe Max-Planck Institute for Ornithology for their support.

